# Harnessing Generative Pre-trained Transformer for Antimicrobial Peptide Generation and MIC Prediction with Contrastive Learning

**DOI:** 10.1101/2025.03.07.642021

**Authors:** Keer Hu, Yang Xiao, Xiao Liu, Shaohua Ma

## Abstract

Antimicrobial peptides (AMPs) have garnered considerable attention due to their reduced likelihood of inducing resistance in pathogens compared to traditional antibiotics, which has spurred the interest in *de novo* design of AMPs. Despite the availability of various methods, accurately generating AMPs and predicting their inhibitory effects remains a challenging task. In this work, we introduce AMPCLGPT, a novel approach that leverages contrastive learning and generative pre-training for AMP design and minimum inhibitory concentration (MIC) prediction. First, AMPCLGPT is pre-trained on a large-scale unlabeled peptide dataset to learn peptide sequence patterns and enhance its ability to extract powerful representations. Second, the pre-trained AMPCLGPT is fine-tuned on AMP data with contrastive learning to increase the distance between antimicrobial and non-antimicrobial peptides in the latent space, improving its ability to accurately generate AMPs. Additionally, the pre-trained AMPCLGPT is fine-tuned to predict MIC values based on the learned peptide features. Empirical results demonstrate that our model can effectively generate AMPs and accurately predict their MIC values. By integrating these two capabilities, AMPCLGPT enables fully automated design of AMPs with low MIC values. AMPCLGPT represents a significant advancement in the field of AMP research, potentially accelerating the development of potent AMP-based therapeutics.

## I. Introduction

Antimicrobial peptides (AMPs) are a diverse class of naturally occurring, small, cationic, and amphipathic molecules that serve as crucial components of the innate immune system in various organisms, including humans, animals, and plants [1], [2]. These peptides typically contain 12 to 50 amino acids and exhibit a broad spectrum of antimicrobial activity against bacteria, fungi and viruses. AMPs have garnered considerable attention as promising alternatives to conventional antibiotics due to their unique mechanisms of action, which minimize the development of resistance [3]. Consequently, the study of AMPs has become a rapidly growing area of research in the healthcare and pharmaceutical industries.

Despite the potential benefits of AMPs, their widespread application as therapeutics is hindered by several challenges. First, the identification and isolation of novel AMPs from natural sources is time-consuming and resource-intensive [4]. Second, AMPs often exhibit low bioavailability, rapid degradation, and potential toxicity, which limits their clinical applicability [5]. Therefore, there is a pressing need for innovative approaches to discover, design, and optimize AMPs with enhanced potency, selectivity, and pharmacokinetic properties. Artificial intelligence (AI)-based computational methods have emerged as valuable tools to accelerate and streamline the development of novel AMPs [6], [7].

Recently, AI has made significant advancements and has been increasingly applied to various domains within the life sciences, including drug discovery, genomics, proteomics, and medical imaging [8], [9]. In terms of peptide research, AI has emerged as a valuable tool for the discovery, design, and optimization of novel peptides with desired properties and functions. Computational methods, such as machine learning and deep learning algorithms, can efficiently predict specific biological activities, physicochemical properties, and structural features of AMPs [10], [11]. Furthermore, AI-based techniques can facilitate the optimization of AMP sequences by predicting their stability, solubility, and potential toxicity, which are crucial factors for their therapeutic applicability [12]. By leveraging AI approaches, researchers can overcome the limitations of traditional experimental methods, reduce the time and cost associated with AMP development, and ultimately enhance the success rate of AMP-based therapeutics.

More recently, several approaches have emerged for *de novo* design of AMPs with promising results. These methods primarily employ deep learning techniques, such as Variational Autoencoders (VAEs) and Generative Adversarial Networks (GANs), for generating novel AMP sequences with desired properties [10]–[12]. However, despite the advancements made in these AI-driven approaches, certain limitations remain: 1) Low success rate in AMP design: The current models may generate a considerable number of false-positive samples during the design process, leading to a lower success rate in identifying potent AMPs. This issue can be attributed to the challenges in accurately capturing the complex sequence-activity relationships of AMPs, as well as the inherent biases and noise present in the training data. 2) Low antimicrobial efficacy: The AMPs generated by these methods often exhibit low antimicrobial activity, as indicated by their relatively high minimum inhibitory concentration (MIC) values. This limitation may arise due to the lack of incorporation of advanced optimization techniques or the inability of the models to fully capture the diverse mechanisms of action and structural features that contribute to the potency of AMPs. To address these limitations, it is essential to explore and integrate novel computational techniques for improving the performance of AI-driven methods in AMP design.

Generative Pre-trained Transformers (GPTs) are a class of advanced deep learning models that have demonstrated remarkable success in various natural language processing (NLP) tasks, such as text generation, translation, and sentiment analysis [13], [14]. The success of GPTs in NLP has inspired their application to biological sequence data, which share several characteristics with natural language, such as discrete alphabets (amino acids or nucleotides) and sequential organization. Recent studies have demonstrated the potential of GPTs in various bioinformatics tasks, including DNA analysis and single-cell analysis [15]–[17]. Thus, GPTs hold great promise for advancing peptide research, particularly in the design and optimization of AMPs. By applying GPTs to AMP research, a crucial advantage over previous generative models is the ability to pre-train GPTs on large-scale peptide datasets, followed by fine-tuning on specific AMP data. The pre-training step improves the model’s generalization ability, allowing it to capture the underlying structure and organization of peptide sequences more effectively. This broad understanding of peptide sequence space can lead to the generation of novel and diverse AMP candidates. The fine-tuning step focuses on the specific task of AMP generation, tailoring the model to the specificity of antimicrobial activity. This process enhances the model’s ability to identify potent AMP candidates with desired properties. Despite the considerable potential of GPTs in AMP research, there remains a challenge in distinguishing between antimicrobial peptides and non-antimicrobial peptides after the fine-tuning process. The model may generate peptide sequences that exhibit insufficient discrimination between the two classes, resulting in a lower success rate in identifying potent AMP candidates. To address this issue, it is essential to guide the network to better differentiate between antimicrobial and non-antimicrobial peptides in the latent space, thereby increasing the separation between the two classes. Contrastive learning is a powerful technique that aims to learn meaningful representations by maximizing the similarity between instances of the same class while minimizing the similarity between instances of different classes [18]. By incorporating contrastive learning into the GPT-based framework for AMP generation, the model can be guided to enhance the distinction between antimicrobial and non-antimicrobial peptides in the latent space. This approach can lead to improved discrimination between the two classes during the generation process, ultimately resulting in a higher success rate in identifying potent AMP candidates.

Based on the observations, we propose a novel approach called AMPCLGPT (**A**nti**m**icrobial **P**eptide generation with **C**ontrastive **L**earning using **G**enerative **P**re-trained **T**ransformers) for the generation of antimicrobial peptides and the prediction of their minimum inhibitory concentration (MIC) values. First, the GPT model is initially pre-trained on a large-scale peptide dataset, such as the PeptideAtlas, to learn general sequence patterns and features across diverse peptides. This pre-training step enhances the model’s feature extraction capabilities, allowing it to capture the underlying structure and organization of peptide sequences, which can be beneficial for downstream tasks. Following pre-training, the model is fine-tuned on a dataset of antimicrobial peptides using contrastive learning. This process enables the model to focus on learning the specific characteristics of AMPs and to increase the separation between positive (antimicrobial) and negative (non-antimicrobial) samples in the latent space. By maximizing the similarity within the same class and minimizing the similarity between different classes, the model can generate peptide sequences with improved discrimination between antimicrobial and non-antimicrobial properties. Besides, the pre-trained and fine-tuned model is further finetuned using MIC data to learn how to score and evaluate the antimicrobial potency of a given peptide sequence. This step allows the model to predict MIC values for generated AMP candidates, providing a quantitative assessment of their antimicrobial efficacy. By integrating GPTs, contrastive learning, and a multi-stage fine-tuning process, the AMPCLGPT approach aims to overcome the current limitations in AMP design and optimization, leading to the generation of potent antimicrobial peptides and accurate MIC predictions.

## II. Related Work

### A. Generative Models in AMP Generation

The innovative approach of employing generative models to engineer AMPs has emerged as a pivotal strategy in AMP design. Generative models, particularly those leveraging deep learning architectures, have demonstrated an unparalleled capacity to generate novel peptide sequences with potent antimicrobial properties [10]–[12]. The application of generative adversarial networks (GANs) and variational autoencoders (VAEs) in *de novo* peptide design has opened new avenues for the discovery of bioactive molecules [12], [19]. These models are trained on large datasets of known AMPs and are capable of capturing the essential features that contribute to their antimicrobial efficacy. By understanding the underlying sequence patterns and physicochemical properties, generative models facilitate the creation of peptides with enhanced stability, activity, and reduced toxicity.

### B. GPT in Bioinformatics and Biomedicine

The success of Generative Pre-trained Transformers (GPTs) in natural language processing has inspired their application in the fields of bioinformatics and biomedicine, such as BioGPT [15], DNAGPT [16] and scGPT [17]. BioGPT is a domain-specific generative Transformer language model pre-trained on large scale biomedical literature to generate fluent descriptions for biomedical terms. DNAGPT is a generalized DNA pretraining model trained on over 200 billion base pairs from all mammals for downstream tasks, such as DNA sequence order classifiction and guanine-cytosine content prediction. scGPT is a foundation model for single-cell biology, based on a generative pretrained transformer across a repository of over 33 million cells for cellular biology and genetic research. Despite its success in other domains, the use of GPT for designing AMPs has not been extensively investigated.

### C. Contrastive Learning

Contrastive learning is a powerful machine learning technique that aims to learn meaningful and discriminative representations by maximizing the similarity between instances of the same class (positive pairs) while minimizing the similarity between instances of different classes (negative pairs) [18]. Recently, contrastive learning has be used in various bioinformatics-related applications, particularly in tasks involving high-dimensional and complex biological data [20]– [22]. However, the application of contrastive learning for enhancing AMP generation is not fully explored.

## III. Methods

### A. Overview

Fig. 1 shows the overview of AMPCLGPT, consisting of two stages, *i*.*e*., pre-training and fine-tuning. During the pretraining stage, we utilize a vast amount of unlabeled peptide sequences from the PeptideAtlas database to pre-train the GPT. The objective of this stage is to enable the GPT model to learn the sequence patterns and rules inherent to peptides, which serves as a foundation for the subsequent fine-tuning tasks. The following fine-tuning stage is further divided into two tasks: AMP generation and prediction of MIC values. For the AMP generation task, the pre-trained GPT model is fine-tuned using contrastive learning to enhance the distance between antimicrobial and non-antimicrobial peptides in the latent space. This process aims to guide the model towards generating peptide sequences with potent antimicrobial properties, thereby improving the discrimination between antimicrobial and non-antimicrobial peptides. For the MIC value prediction task, the pre-trained GPT is used to extract features from AMP sequences, which are then fed into a regression neurak network to predict their MIC values. The data examples of different tasks and special tokens are displayed in Fig. 2.

**Fig. 1.**
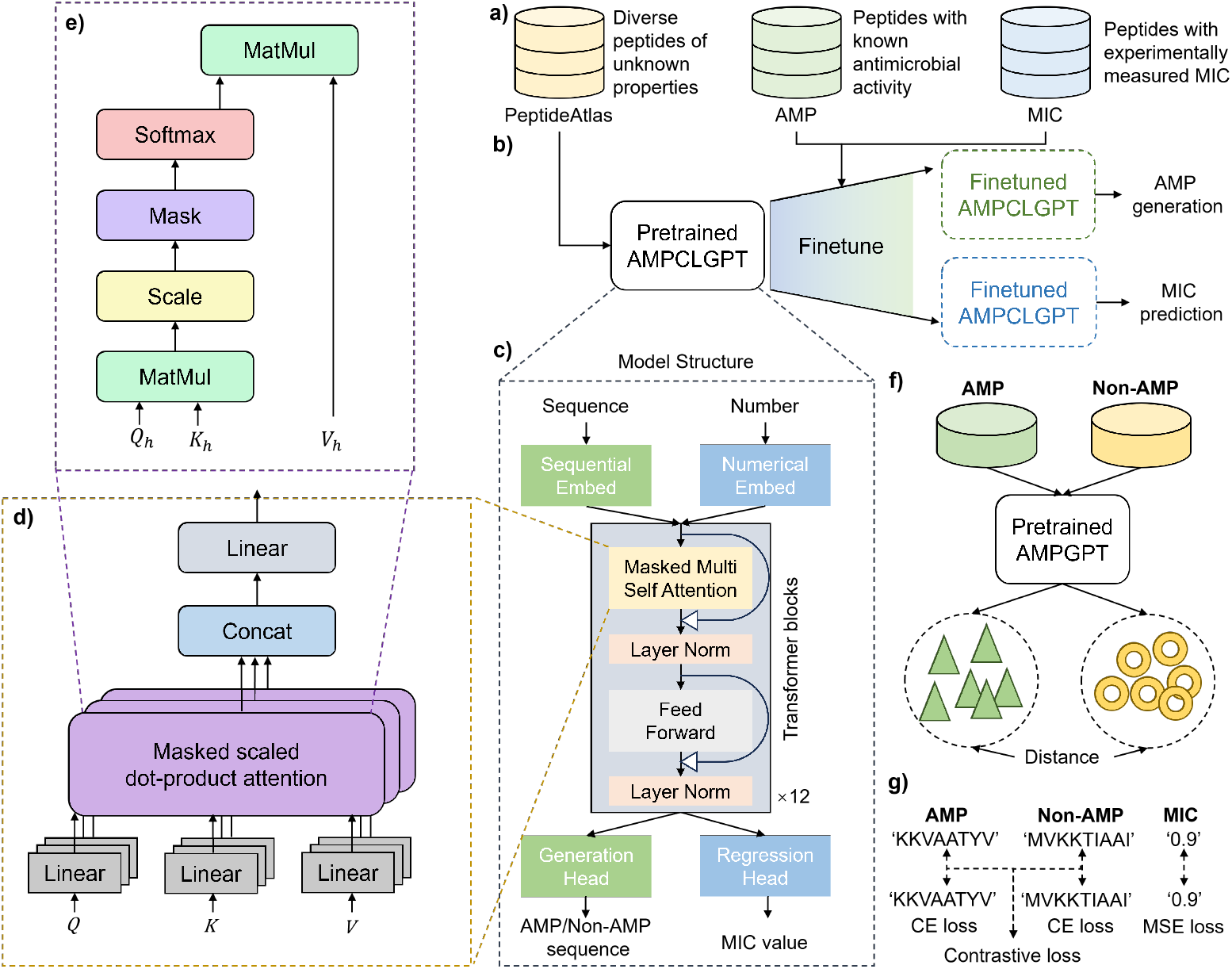
The overall framework of AMPCLGPT.

**Fig. 2.**
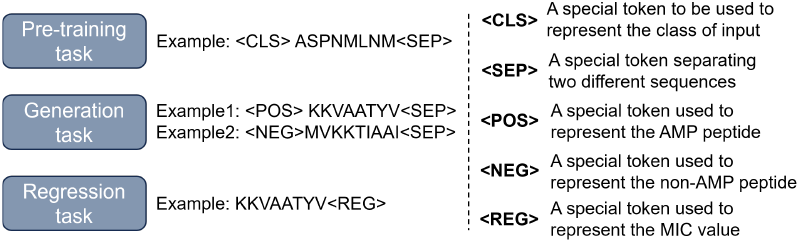
Data examples of different tasks and special token characterization.

### B. Materials and Pre-processing

#### 1) Pre-training data

The pre-training data is collected from PeptideAtlas [23], which is grouped at 50% sequence identity with MMSeqs2 [24] to ensure adequate sequence variation between the training and validation sets. The final training set contains 984,013 sequences and the validation set contains 69,123 sequences.

#### 2) AMP data

The AMP dataset is composed of both positive and negative examples. Positive examples are derived from manually curated AMP databases. We integrate experimentally validated peptides from the dbAMP [25], AMP Scanner [26], and DRAMP [27] databases, ensuring duplicates are eliminated. Negative examples are treated as biologically inactive and are sourced manually using UniProt search filters, specifying subcellular location as cytoplasm and excluding properties such as antimicrobial, antibiotic, antiviral, antifungal, effector, and excreted. To enhance dataset diversity, negative sequences with ≥ 40% sequence identity are removed and substituted with representative sequences from clusters using MMseqs2 [24]. The final set of representative sequences only includes those shorter than or equal to 25 amino acids, and, to prevent length bias, negative sequences are randomly truncated to fit the positive dataset’s length distribution. Both the positive and negative datasets encompass 11,131 sequences each.

#### 3) MIC data

The MIC dataset includes peptides with experimentally confirmed antimicrobial activity and recorded MIC values from the GRAMPA [28] database. Out of 8049 entries, only 4546 are unique sequences. We focus on those peptides that have been experimentally tested for activity against *E. coli* strains. For peptides with multiple measurements, MIC values are averaged, resulting in 4546 entries. Only 3444 peptides meet the required length of 25 amino acids.Peptides are labeled as active (positive) if their MIC is*<*=10^1.5^(32 µg/ml), while those with MIC *>*32 µg/ml are considered inactive (negative).

### C. AMPCLGPT

#### 1) Model Architecture

Fig. 1(c) illustrates the structure of our proposed AMPCLGPT, which is composed of a stacked of Transformer blocks. Each Transformer block consists of a multi-head self-attention (MHSA) module, followed by position-wise feed-forward layers. These components are interconnected through residual connections and layer normalization, enabling the model to effectively learn complex sequence patterns and dependencies.

The input to the model is a peptide sequence, represented as a sequence of discrete tokens corresponding to the amino acids. The tokens are first embedded into a continuous vector space, and positional encodings are added to capture the sequential nature of the input data. The resulting embeddings are then fed into the stack of Transformer blocks, where the self-attention mechanism allows the model to capture both local and long-range dependencies within the peptide sequence. As the model processes the input through the Transformer blocks, it learns to generate contextually relevant peptide sequences in an autoregressive manner. During the fine-tuning stage for AMP generation, the model is optimized using contrastive learning to enhance the separation between antimicrobial and non-antimicrobial peptides in the latent space. For the MIC value prediction task, the model extracts features from the peptide sequences, which are then passed through a regression layer to predict the corresponding MIC values.

#### 2) MHSA module

The MHSA module is the crucial component of the Transformer block, as illustreated in Fig. 1(d). The MHSA module allows the model to focus on different parts of the input sequence for each attention head, thereby capturing various aspects of the sequence information.

Given a sequence of input embeddings 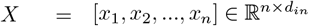, where *x*_*i*_ represents the embedding of the *i*-th token in the sequence, *d*_*in*_ is the dimension of the input token, and *n* is the sequence length, the MHSA module first projects *X* into three different spaces to obtain the query (*Q*), key (*K*), and value (*V* ) matrices:

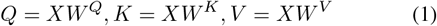

where 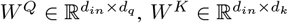, and 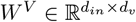 are learnable weight matrices. The projected queries, keys and values are of size *d*_*q*_, *d*_*k*_ and *d*_*v*_, respectively. This allows each head to operate on different and learnable linear projections of the input embedding. To control the size of each head, we following the previous study to use the policy: *d*_*q*_ = *d*_*k*_ = *d*_*v*_. To incorporate multiple attention heads, the model is extended as follows:

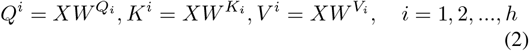

where *h* is the number of attention heads and 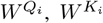 are the learnable weight matrices for the *i*-th attention head. Subsequently, the attention scores between the query and key pairs for each head are computed as shown in Fig.1(e):

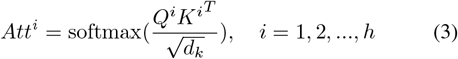

where the square root operation in the denominator serves as a scaling factor to prevent the dot product from growing too large. The output of the MHSA module for each head is obtained by weighting the value vectors *V* ^*i*^ with the attention scores *Att*^*i*^:

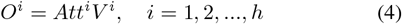

Finally, the outputs from all heads are concatenated and linearly transformed using a learnable weight matrix 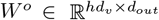 to produce the final output of the MHSA module:

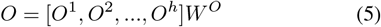

#### 3) Pre-training to Learn Peptide Distribution

The pretraining stage is a crucial step in our proposed method, as it aims to learn the underlying distribution of peptide sequences from a large-scale dataset. To achieve this, we employ the AutoRegressive Language Modeling (ARLM) objective, which is a widely used pre-training strategy for GPT and other autoregressive Transformer-based models. In ARLM, the model is trained to predict each token in the sequence based on the context provided by the preceding tokens.

Given a peptide sequence *X* = [*x*_1_, *x*_2_, …, *x*_*n*_], the model processes the input sequence in an autoregressive manner and generates a probability distribution over the vocabulary for each position:

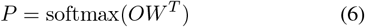

where 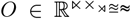 is the output of the MHSA module, 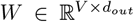 is a learnable weight matrix, and *V* is the size of the vocabulary. The model is trained to minimize the negative log-likelihood of the tokens in the sequence:

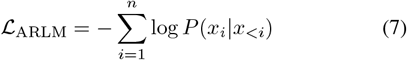

where *x*_*<i*_ denotes the context of all tokens preceding the *i*-th token in the sequence. This pre-training step improves the model’s generalization ability, allowing it to effectively learn the sequence patterns and features that are crucial for the subsequent fine-tuning tasks.

#### 4) Fine-tuning to Generate AMP with Contrastive Learning

Following the pre-training stage, the model is fine-tuned for the task of AMP generation using contrastive learning. This stage aims to enhance the model’s ability to differentiate between antimicrobial and non-antimicrobial peptides and to generate potent AMP candidates.

Formally, given a pair of peptide sequences (*X, Y* ), the model computes their representations *f* (*X*) and *f* (*Y* ), and the distance between these representations is measured using the KL divergence:

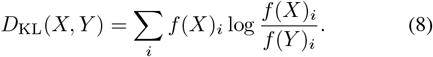

Then the contrastive loss function is defined as:

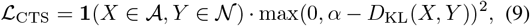

where 𝒜 represents the set of antimicrobial peptides 𝒩, represents the set of non-antimicrobial peptides, **1**(·) is the indicator function, and *α* is a margin parameter that controls the distance between the two classes in the latent space. The contrastive loss is combined with the autoregressive loss that maintains the model’s ability to generate coherent AMP sequences. The joint loss function for the fine-tuning stage is then defined as a weighted sum of the contrastive loss and the autoregressive loss:

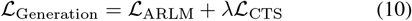

where *λ* is a hyper-parameter that controls the trade-off between the two losses.

#### 5) Fine-tuning to Predict MIC of AMP

The pre-trained model is also fine-tuned for the task of predicting the MIC values of AMPs. This task provides a quantitative assessment of the antimicrobial efficacy of the generated peptides, thereby facilitating the identification of potent AMP candidates. Given a peptide sequence *X*, the model computes its representation *f* (*X*), followed by a regression layer to predict the MIC value:

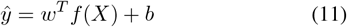

where *w* is a learnable weight vector and *b* is a bias term. The model is trained to minimize the mean squared error (MSE) between the predicted MIC values and the true MIC values:

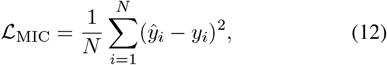

where *N* is the number of samples in the dataset, *ŷ*_*i*_ and *y*_*i*_ are the predicted and true MIC values for the *i*-th AMP, respectively.

#### 6) Peptide Generation

For the peptide generation process, we employed a top-k sampling strategy to ensure the diversity and quality of the generated antimicrobial peptides (AMPs). In our implementation, we set k = 10.

## VI. Experiments AND Results

### A. Implementation Details

All experiments are conducted on the same machine with Intel(R)Core(TM)i9-7900X CPU @ 3.30GHz and 4 GPUs (NVIDIA TESLA V100). For model evaluation, we split the pre-training and fine-tuning dataset into training, validation and test sets with a 7:1:2 ratio, respectively.

### B. AMPCLGPT can Effectively Generate AMPs that Are Novel with Desired Properties

As illustrated in the fig3 and fig4, our AMPCLGPT model successfully generates peptide sequences with an amino acid composition closely resembling that of known antimicrobial peptides (AMPs). Notably, the model maintains a consistent preference for the most frequently occurring amino acids such as glycine (G), lysine (K), and leucine (L), as well as the least frequent residues like histidine (H) and methionine (M). This suggests that the model has effectively learned the characteristic amino acid distribution patterns inherent to AMPs. Furthermore, our model is capable of producing peptides with desired physicochemical properties. We examined the generated peptides for key properties including aromaticity, charge, hydrophobicity, hydrophobic moment, and length. According to previous studies,distribution comparison reveals that the positively charged peptides exhibit significantly higher charge, hydrophobic moment, and hydrophobicity than the negative counterparts, as evidenced by the one-sided Mann-Whitney U test (p-value*<*0.05) [12]. The results indicate that the physicochemical profiles of the generated peptides closely align with those of real AMPs. our model has successfully captured certain intrinsic properties that are critical to the antimicrobial activity of AMPs. Moreover, Fig 5 presents the minimum edit distance between the peptides generated by our model and those in the known AMP databases. The results indicate that our model does not merely replicate existing AMP sequences; instead, it has learned to explore a broader antimicrobial peptide space. This suggests that AMPCLGPT is capable of generating novel peptide sequences that diver from known templates, potentially uncovering new regions of the sequence space that may lead to the discovery of previously unidentified antimicrobial agents.

**Fig. 3.**
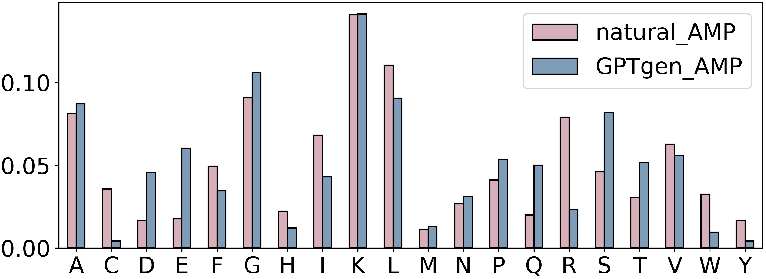
Comparison of amino acid composition between generated AMP by our AMPCLGPT and true AMPs.

**Fig. 4.**
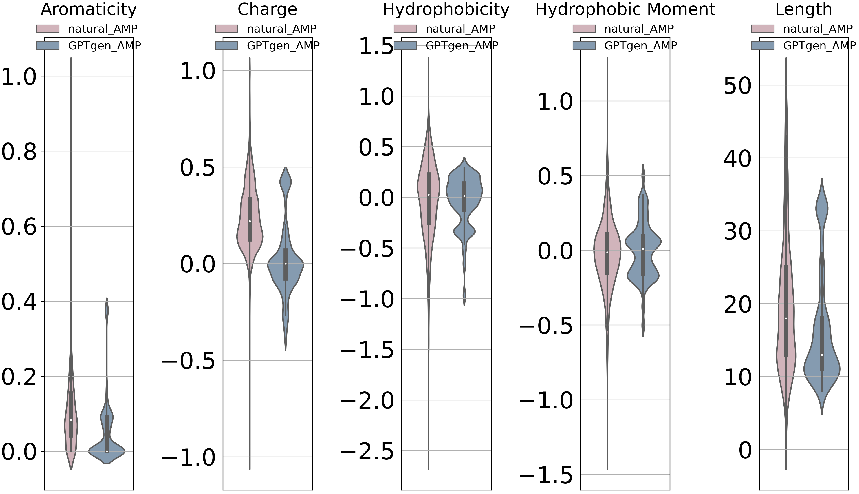
Comparison of key molecular features between generated AMP by our AMPCLGPT and true AMPs. From left to right: Aromaticity, Charge, Hydrophobicity, Hydrophobic Moment, and Length.

**Fig. 5.**
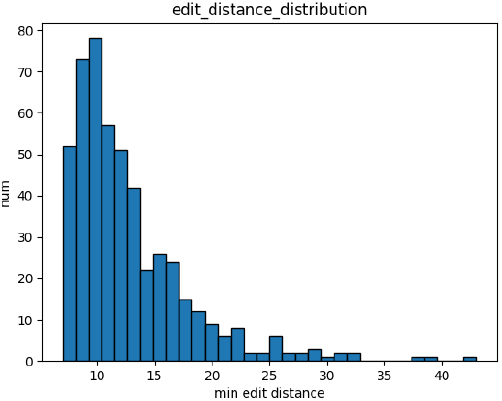
Comparison of amino acid composition between generated AMP by our AMPCLGPT and true AMPs.

### C. AMPCLGPT can Accurately Predict AMP’s MIC

Predicting the minimum inhibitory concentration (MIC) of AMPs can be formulated as a regression task. To address this, we designed a regression head within our AMPCLGPT framework specifically for MIC prediction. We conducted a comparative analysis against several prevalent machine learning models, including Random Forest (RF), Gradient Boosting (GB), and XGBoost, as well as commonly used deep learning models like Convolutional Neural Networks (CNNs). Additionally, we benchmarked our model against a modified Long Short-Term Memory (LSTM) model from the literature [29].

As shown in Table 1, our model achieved a Pearson correlation coefficient of 0.61 between the predicted MIC values and the true MIC values, significantly outperforming the other models. This high correlation indicates that our model provides a more accurate prediction of antimicrobial efficacy, reinforcing the utility of AMPCLGPT as a robust tool for guiding the design of AMPs with desired inhibitory properties.

**TABLE I.** Comparison BETWEEN OUR METHOD AND BASELINE METHODS FOR MIC PREDICTION.

| Method | Type | Pearson |
| --- | --- | --- |
| RF | Machine Learning | 0.2375 |
| GB | Machine Learning | 0.3162 |
| XGBoost | Machine Learning | 0.4202 |
| AMPCNN | Deep Learning | 0.4045 |
| AMPLSTM | Deep Learning | 0.5098 |
| AMPCLGPT (w/o pre-training) | Deep Learning | 0.4907 |
| AMPCLGPT | Deep Learning | **0.6137** |

### D. Utilizing AMPCLGPT to Generate AMP with Low MIC

By utilizing our MIC regression head as a predictive tool, we can effectively identify candidate peptides with lower MIC values, thereby reducing the experimental workload in wetlab settings. As depicted in Fig6 and Fig7, the amino acid composition and physicochemical properties of the peptides we selected with the lowest predicted MIC values closely mirror those of the known AMPs with the lowest MIC values in the real data (sample size = 25). This consistency underscores the capability of our model to not only predict MIC values with high accuracy but also to select peptides that share key characteristics with highly potent AMPs. And in Figure 8, we found significant differences in some physicochemical properties (aromaticity and charge) between peptides with lower MIC values predicted by our model and peptides with lower MIC values, suggesting that higher or lower levels of these two physicochemical properties may be associated with higher MIC values.Antimicrobial peptides were also shown to be significantly higher than non-antimicrobial peptides in these two metrics in a previous study [12]. In addition, we employed the AlphaFold3 tool to simulate the structures of both real and generated peptides with low MIC values [30]. The simulations revealed that the majority of the peptides generated by our model exhibit high predicted local distance difference test (pLDDT) scores, indicating a high confidence in the predicted structures. This suggests that our model has effectively learned the natural sequence patterns that govern peptide folding and stability. Moreover, both the real and generated low MIC value peptides predominantly exhibit alpha-helical structures. The presence of alpha helices is known to correlate with the stability of peptides. Interestingly, we observed the occurrence of slight to moderate unstructured regions at the ends of the peptide chains. These unstructured regions may play a role in the mechanism by which antimicrobial peptides penetrate and integrate into cell membranes, a key aspect of their function [12]. These structural insights not only support the validity of our model’s predictions but also enhance our understanding of the functional properties of the generated peptides.

**Fig. 6.**
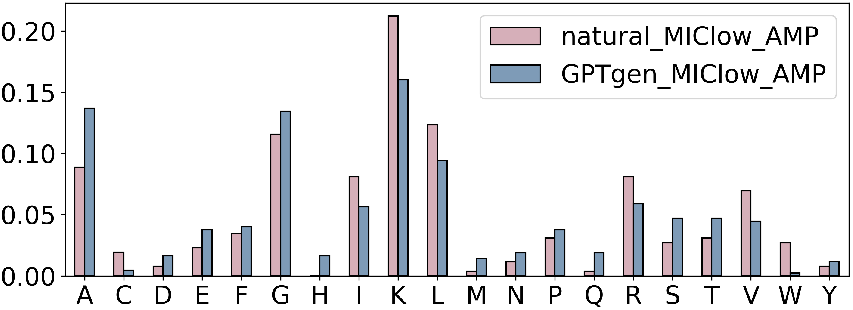
Comparison of amino acid composition between generated AMPs with low MIC values filtered by AMPCLGPT and true low-MIC AMPs.

**Fig. 7.**
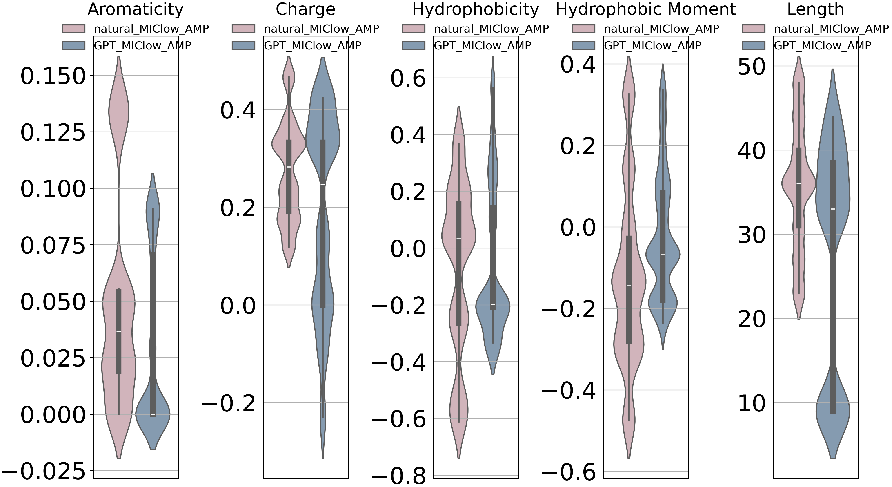
Comparison of key molecular features between generated AMPs with low MIC values filtered by AMPCLGPT and true low-MIC AMPs.

**Fig. 8.**
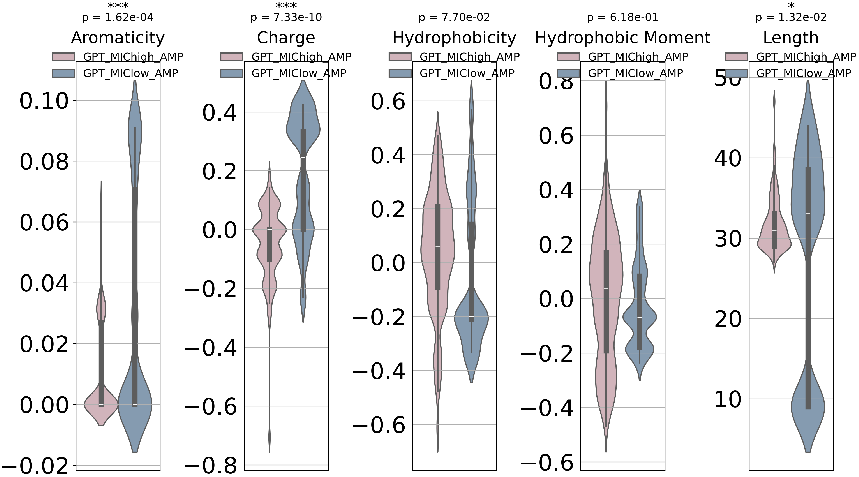
Comparison of key molecular features between generated AMP with low MIC values and high MIC values by our AMPCLGPT.

**Fig. 9.**
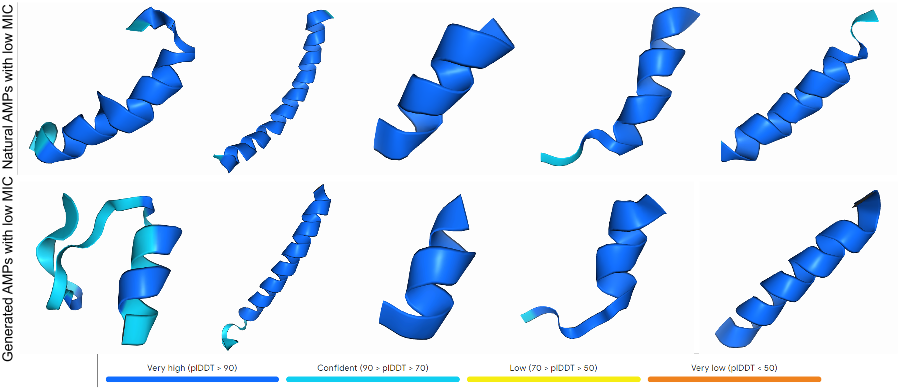
Structures of generated AMPs and true AMPs with low MIC values.

### E. Ablation Study

#### 1) Ablation Study on the Generation Task

In our generation experiments, we employed cross-entropy (CE) as the metric to evaluate the training effectiveness of the GPT model, where a lower cross-entropy value indicates better performance. As shown in Table 2, when either the pre-training module or the contrastive learning module is removed, the cross-entropy value increases to some extent. This also shows the significance of our pre-training in large peptide libraries and the effectiveness of adding a comparative learning module to differentiate between generating antimicrobial and non-antimicrobial peptides

**TABLE II.** Ablation STUDY OF THE PROPOSED METHOD FOR AMP GENERATION TASK. (CE: Cross Entropy)

|  | CE (Training set) | CE (Test set) |
| --- | --- | --- |
| w/o pre-training | 3.6051 | 3.6017 |
| w/o contrastive learning | 3.6048 | 3.6122 |
| AMPCLGPT | 2.7044 | 2.6988 |
| Uniform random | 25 | 25 |

#### 2) Ablation Study on the Regression Task

In the regression task’s ablation study, we removed the pre-trained weights to evaluate their impact on the model’s performance. The results showed a decrease in the Pearson correlation coefficient (R value), indicating a decline in the model’s ability to accurately predict MIC values. This decline underscores the critical role of the pre-training module in enhancing the model’s performance.

## V. Discussion

There are many hyperparameters that can be explored during the sampling process, such as the k parameter in topk. Sampling generates creative an be degenerate or not sample natural sequence propensities for small values of k. Larger values of k produce sequences whose propensities match natural ones. More importantly, to further enhance the effectiveness and applicability of AMPCLGPT, several key areas warrant improvement. First, while AMPCLGPT has shown impressive results in generating and predicting antimicrobial peptides (AMPs) with low minimum inhibitory concentration (MIC) values, it is crucial to validate these computationally designed peptides through wet-lab experiments. By synthesizing the generated peptides and testing their antimicrobial activity against various bacterial strains, the practical applicability and effectiveness of AMPCLGPT’s outputs can be confirmed. Such validation would not only strengthen the model’s credibility but also refine its predictions by providing feedback to improve its underlying algorithms.

## VI. Conclusion

In this study, we introduced AMPCLGPT, a novel approach that leverages Generative Pre-trained Transformers (GPT) and contrastive learning for the generation of antimicrobial peptides (AMPs) and the prediction of their minimum inhibitory concentration (MIC) values. Through extensive experimentation, we demonstrated that AMPCLGPT is capable of generating peptides with amino acid compositions and physicochemical properties that closely resemble those of known AMPs. The model’s ability to generate novel sequences that maintain the critical features of effective AMPs while exploring a broader peptide space highlights its potential in advancing the discovery of new antimicrobial agents.

